# Structural determinants of Acid-sensing ion channel potentiation by single chain lipids

**DOI:** 10.1101/2021.06.25.449979

**Authors:** Robert C. Klipp, John R. Bankston

## Abstract

Acid-sensing ion channels (ASICs) are sensitized to activation by inflammatory mediators like the polyunsaturated fatty acid (PUFA) arachidonic acid (AA). Previous work has shown that AA can potentiate ASIC currents at subsaturating proton concentrations, but the structural mechanisms of this change in gating are not understood. Here we show that PUFAs cause multiple gating changes in ASIC3 including shifting the pH dependence of activation, slowing the rate of desensitization, and increasing the current even at a saturating pH. The impact on gating depends on the nature of both the head and tail of the lipid with the head group structure primarily determining the magnitude of the effect on the channel. An N-acyl amino acid (NAAA), arachidonoyl glycine (AG), is such a strong regulator that it can act as a ligand at neutral pH. Mutation of an arginine in the outer segment of TM1 (R64) eliminated the effect of docosahexaenoic acid (DHA) even at high concentrations suggesting a potential interaction site for the lipid on the channel. Our results suggest a model where PUFAs bind to ASICs both via their tail group as well as an electrostatic interaction between the negatively charged PUFA head group and the positively charged arginine side chain. These data provide the first look at the structural features of lipids that are important for modulating ASICs and suggest a potential binding site for PUFAs on the channel.

**Short Summary:** Arachidonic acid (AA) is a known modulator of ASICs. Bankston and Klipp examine the structural requirements for ASIC modulation by a number of lipids related to AA. Negatively charged head groups are stronger potentiators and may interact with an arginine in TM1 near the outer leaflet of the plasma membrane.

## Introduction

Acid-sensing ion channels (ASICs) are voltage-insensitive, pH activated members of the Deg/ENaC family of ion channels^1^. ASICs are widely expressed throughout the body including in neurons in both the central (CNS) and peripheral nervous system (PNS)^1^. They have been demonstrated to impact the sensing of pain and inflammation in the PNS as well as fear conditioning, ischemic cell death, and synaptic plasticity in the CNS^1^. The physiological activation mechanisms of ASICs are not well understood. In pathophysiological conditions like stroke, neurons are exposed to prolonged acidosis^2^. During normal synaptic transmission, it is hypothesized that the synapse becomes transiently acidic after vesicular release of neurotransmitter because the vesicles have a pH ~5^3^. However, this transient, and usually modest, acidification is unlikely to activate many channels in most synapses. There is a growing body of literature suggesting that several endogenous molecules including serotonin, lactate, and nitric oxide act to sensitize ASICs to protons making them more active at less acidic pH^4–7^.

The lipid composition of the membrane can significantly impact the function of ion channels. KCNQ1 and GIRK channels require phosphatidylinositol 4,5-bisphosphate (PIP2) to function while cholesterol can alter the function of nAChR and TRPV1 channels^8–10^. Polyunsaturated fatty acids (PUFAs) have been shown to modulate the large-conductance Ca^2+^- and voltage-gated K^+^ (Slo1 BK) channel with an EC_50_ in the hundreds of nanomolar^11–13^. In addition, the voltage-gated potassium channel, K_V_7.1, can be either potentiated or inhibited by PUFAs and PUFA-derivatives depending on the structure of the fatty acid^14–20^.

Overall, little is known about the role lipids play in ASIC function. A recent structure of full-length ASIC1 from chicken extracted from membranes using SMA copolymer, solved at 2.8Å resolution, revealed multiple ordered elongated densities consistent with lipids interacting along the transmembrane domains of the channel^21^. Previous structures in detergent micelles adopted substantially different conformations suggesting that lipids are vital to ASIC structure and function^22–24^. A small number of papers have looked at regulation of ASICs by the PUFA arachidonic acid (AA)^25–28^. These studies have shown that AA potentiates ASIC1a and ASIC3 currents likely by shifting the pH dependence of channel activation through a direct action on the channel^25,26^. This effect leads to increased pH-activated currents in DRG neurons as well as increased action potential firing in response to modest acidification of the extracellular space around the neuron^26,27^. In addition, exudates, containing lysophophatidylcholine (LPC) and AA, from human patients with inflamed joints could activate ASICs without any change in pH^28^. These early studies suggest that lipids may act as critical sensitizing compounds for this class of channel. However, the structural requirements for lipid action on the channel and a mechanistic hypothesis for how these lipids cause changes in channel function is lacking.

Docosahexaenoic acid (DHA) and AA are particularly abundant in the brain and other neuronal tissues^29^. Unesterified PUFAs can result from either synthesis of dietary precursors like linoleic acid, or through liberation from glycerophospholipids via enzymes like phospholipase A2^30^. PUFAs are nutraceuticals with potential to become lead compounds for rational drug design for targeting ion channels and membrane receptors. For instance, PUFAs are being designed as a potential treatment for patients with long QT syndrome type 1^14,17,19,20^. ASICs may present interesting targets for the inhibition of pain and the inhibition of cell death in both ischemia and neurodegenerative disorders^2,31,32^.

Here, we examine the structural mechanisms of ASIC regulation by PUFAs and PUFA-derivatives. We show that AA potentiates ASIC3 currents by impacting several biophysical properties of the channel. In addition, by looking at more than 30 different PUFAs and PUFA-derivatives we clearly demonstrate that the ability of PUFAs to act on the channel depends on both the head and tail group of the lipid. The properties of the tail group, including length and double bond number, can alter the magnitude of the effect of the PUFA on ASIC3. However, our data show that the head group is likely the critical determinant of lipid efficacy. A head group more prone to harboring a negative charge is more likely to act as a strong potentiator of ASIC3 currents. Replacing the carboxyl head group of PUFAs with a more negatively charged head group leads to significantly larger effects on the channel with some lipids capable of acting as a ligand, activating the channel at neutral pH. Finally, we identified a putative interaction site for the PUFA head group on the first transmembrane segment of ASICs near the outer leaflet of the membrane. Taken together, these experiments provide the first look at the role that lipid structure plays in altering ASIC function as well as hypothesize a potential site on the channel for regulation of function by lipids.

## Results

### PUFAs stabilize the open state of ASICs

AA is a highly abundant PUFA with a 20-carbon tail and four cis double bonds. To indicate the length of the tail and the position of the double bonds for the lipids in this manuscript we will use the following nomenclature, [20(5,8,11,14)], which indicates a 20-carbon tail with four cis double bonds at positions 5, 8, 11 and 14. Previous work has shown that ASIC1a and ASIC3 currents elicited by steps to pH 6.9 are increased upon application of 5μM AA to cells expressing these channels^26^. This increase has been shown to be, at least in part, due to an alkaline shift in the pH dependence of activation. We confirmed this initial observation by applying 10μM AA to Chinese Hamster Ovary (CHO) cells expressing ASIC3 with a C-terminal cerulean tag and measured the pH_0.5_ of activation using the whole-cell patch clamp configuration. Coverslips with cells expressing ASIC3 were incubated in 10μM AA for at least 10 minutes and then the currents were elicited via successive steps to more and more acidic pH (Fig. 1A). We found that 10μM AA shifted the pH_0.5_ of activation by 0.11 pH units yet had no effect on the pH_0.5_ of desensitization, in agreement with previous results (Fig. 1B)^27^.

**Figure 1.**
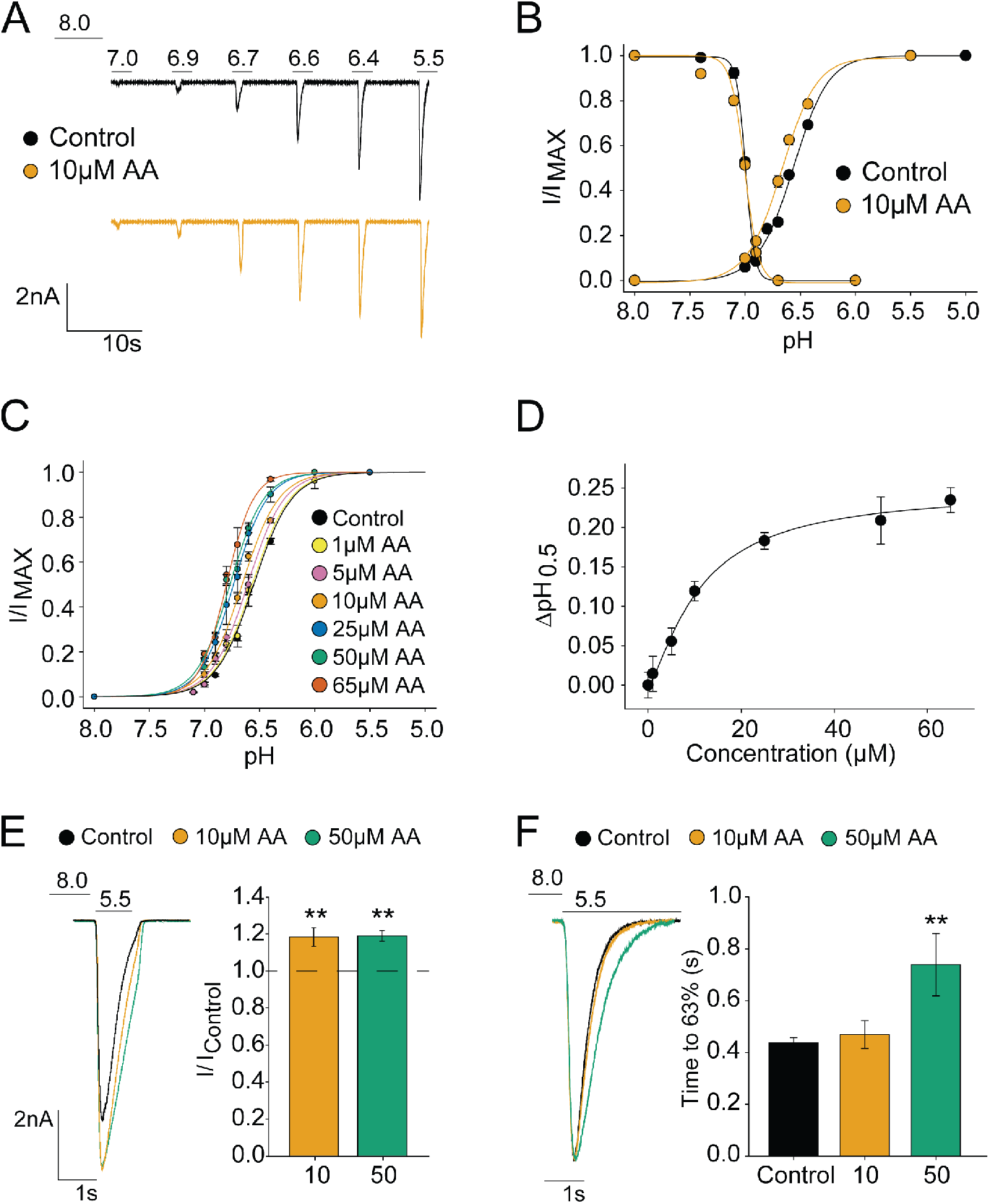
Binding of AA favors the open state of the channel (A) Representative whole-cell recordings showing pH dependent activation of ASIC3 ± 10μM AA. (B) pH activation and desensitization curves for ASIC3 ± 10μM AA (Activation: pH_0.5_ = 6.68 ± 0.01, n = 7 for AA and 6.56 ± 0.01, n = 70 for Control; p = 0.005. Desensitization: pH_0.5_ = 7.00 ± 0.01, hill slope = 7 ± 1.1, n = 6 for AA; and pH_0.5_ = 7.00 ± 0.00, hill slope = 9.8 ± 0.53, n = 5 for Control). (C) Curves showing the pH dependence of activation of ASIC3 at different concentrations of AA. ΔpH_0.5_ values from data in C plotted as a function of concentration yields a half-maximal activation concentration (EC_50_) = 11μM ± 2μM. (E) (left) Representative pH 5.5-evoked currents from a single cell at different concentrations of AA. (right) Bar plot showing the fractional change in the maximum current measured at pH 5.5 (I_MAX_) in response to AA (Fold-increase in I_MAX_ = 1.18 ± 0.05, n = 13 for 10μM AA; and 1.27 ± 0.09, n = 10 for 50μM AA). (F) (left) Representative traces normalized to control peak and (right) bar plot showing the time point at which 63% decay of peak ASIC3 current is reached at different concentrations of AA measured at pH 5.5 (t_63_% = 427ms ± 17ms, n= 19 for Control; 469ms ± 53ms, n = 5 for 10μM AA; and 739ms ± 120ms, n = 5 for 50μM AA). All data given as mean ± SEM. Asterisks in E represent p-values as determined by paired t-tests. Asterisks in F represent p-values as determined by post-hoc Dunnett’s test. ** p < 0.01 (see methods for details).

In order to extend these prior observations, we measured the concentration dependence of the shift in the pH_0.5_ of activation. Plotting the shift as a function of AA concentration and fitting to a Hill-type equation (see methods) yielded an EC_50_ of 11.62μM ± 2.08μM (Fig. 1C,1D). This has not been previously measured for ASICs but compares reasonably to values obtained for several other channels that show EC_50_ values can range between 400nm and 100μM^11,17,33^. The maximum shift in the pH_0.5_ of activation measured at any concentration of AA was 0.23 ± 0.02 pH units.

In addition to the shift in activation pH, we observed two previously unreported changes to ASIC3 gating upon AA application. First, there was an increase in the current magnitude upon addition of AA at saturating proton concentrations (saturating proton concentrations determined from pH_0.5_ curve) (Fig. 1B,C). The amplitude of this current, which we term I_MAX_, was increased ~1.2 fold upon addition of 10μM AA (Fig. 1E). Surprisingly, there was no additional I_MAX_ increase at 50μM AA despite the increased shift in activation pH_0.5_. Second, at a maximally activating pH of 5.5, a concentration dependent slowing of the rate of desensitization was observed upon addition of AA (Fig. 1F). To quantify this result, we measured the time it takes for the current to decrease 1/e of its original value and found that 50μM AA slowed this time from 427.21ms ± 17.75ms to 739.56ms ± 120.58ms (Fig. 1F). Taken together, these data suggest that AA binding favors the open state of the channel.

### Head groups with lower pKa are stronger potentiators of ASIC3 currents

Structurally, PUFAs contain a negatively charged carboxylic acid head group and a long non-polar acyl tail that can have one to six double bonds in varying positions along the tail. Studies in other ion channels have demonstrated that the tail length, number, and position of double bonds, as well as the charge and size of the head group can all contribute to ion channel potentiation^11,16,20,34^. We therefore sought to determine the important structural elements in PUFA modulation of ASICs.

There are AA-derivatives that vary in their head groups that allowed us to first probe the contribution of the head group in ASIC modulation. We compared AA to eight AA-derivatives with differing head groups and identical tails [20(5,8,11,14)] (Fig. 2A) (Table 1). These head groups vary in size, atomic composition, and propensity to ionize with some that readily ionize (e.g., glycine) and others that do not (e.g., methyl ester).

**Figure. 2.**
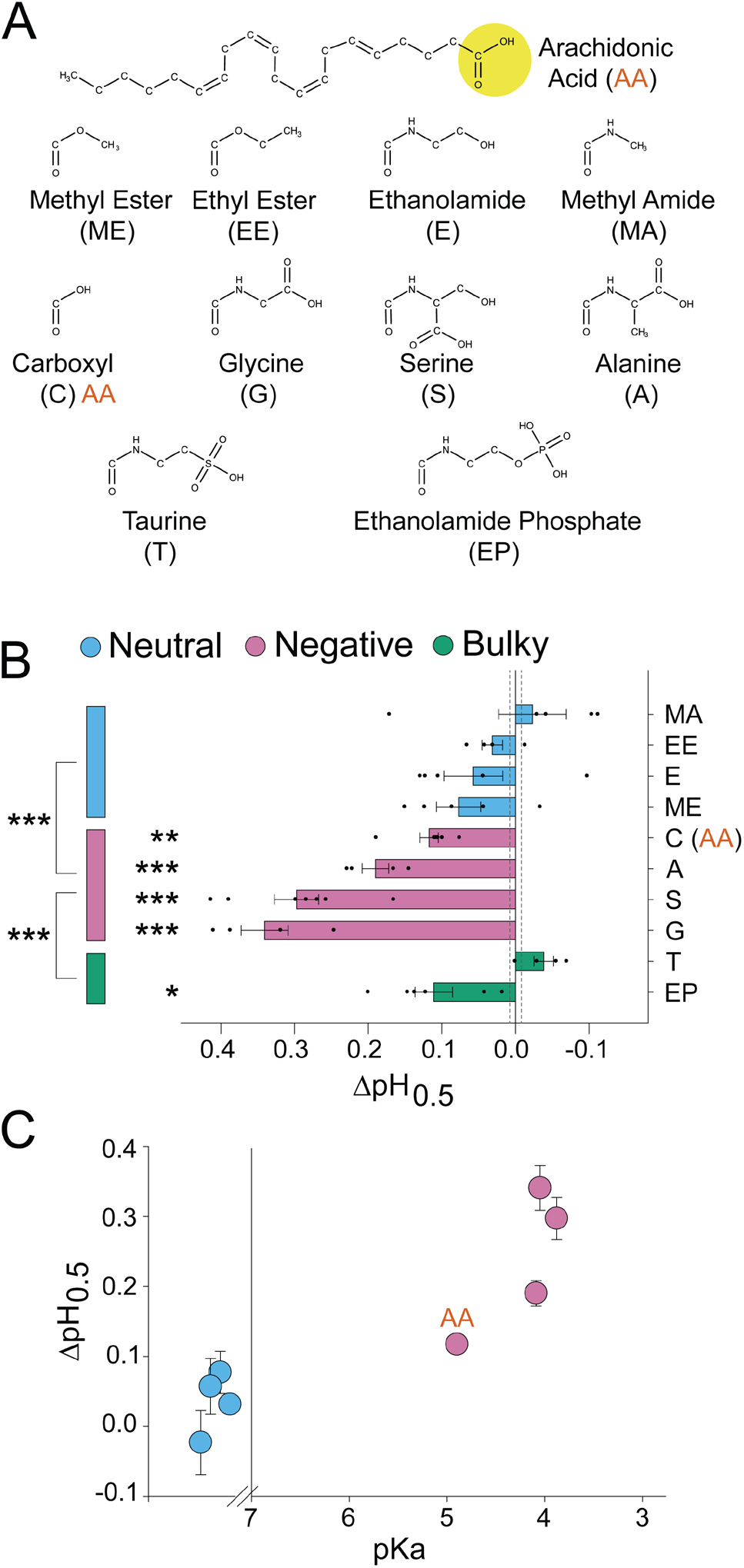
A negatively charged PUFA head group is critical for potentiation of ASIC3. (A) (top) Structures of AA and (below) head groups used to replace AA’s native carboxyl head group, creating the AA-derivatives used in B and C. (B) Activation ΔpH_0.5_ values for ASIC3 in the presence of 10μM of the indicated AA-derivatives from A (n = 4-7, Data given in Table 1). Vertical dashed lines indicate SEM of control (n = 70). (C) ΔpH_0.5_ values from B plotted as a function of the of the calculated pKa for the AA-derivatives. Neutral head groups (blue) are plotted on a non-numbered scale to the left of the break in x-axis (at pKa = 7) representing that they either are permanently neutral, or the pKa is sufficiently large that head groups remain neutral at all pH values tested (calculated pKa values are given in Supplementary Table 2). All data given as mean ± SEM. Asterisks next to horizontal bars in B represent p-values as determined by a post-hoc Dunnett’s test. Asterisks next to vertical bars in B represent p-values as determined by post-hoc Tukey’s test. * p < 0.05, ** p < 0.01, *** p < 0.001 (see methods for details).

**Table 1:**
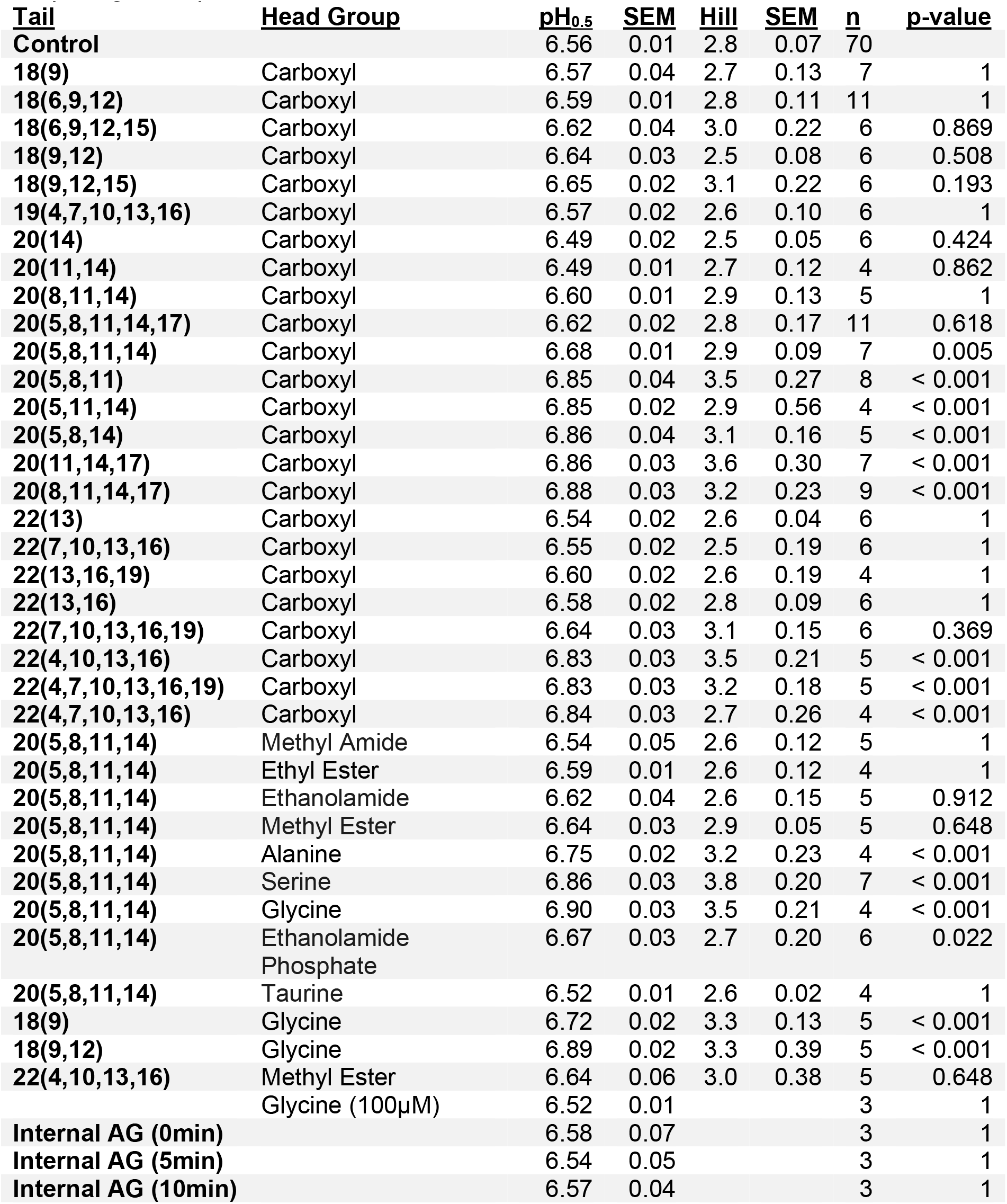
Activation pH_0.5_ values and Hill slopes for ASIC3 in the presence of PUFAs and PUFA-derivatives. All values are measured at 10μM external application of lipid unless otherwise indicated. p-values were calculated with a post-hoc Dunnett’s test comparing each pH_0.5_ dataset to the Control dataset.

To compare potentiation among these AA-derivatives, we again measured the pH dependence of activation in the presence of 10μM lipid (Table 1). Lipids with carboxyl, glycine, serine, or alanine head groups caused significant alkaline shifts in the pH_0.5_ (Fig. 2B). Methyl ester, ethanolamide, methyl amide, and ethyl ester head groups had no significant effect on the pH dependence of activation (Fig. 2B).

In several voltage-gated ion channels, a negatively charged head has been crucial for potentiation^11,16,34^. Figure 2C shows the shifts in the activation pH_0.5_ from figure 2B plotted as a function of the calculated pKa of the AA-derivatives (Calculated values in Supplementary Table 2). The neutral head groups are shown to the left of the scale break on the x-axis at pKa = 7, representing either their lack of ionizable atoms (ethyl ester and methyl ester), or a pKa well outside the range of pH values used in our experiments (ethanolamide and methyl amide). In general, head groups with a lower pKa tended to have a larger potentiation of ASIC3 currents. The reported pKa values were calculated using MarvinSketch and reflect the pKa of the head group connected to the acyl tail in solution, not in a bilayer. Certainly, the local environment of the lipid will have a large impact on the charge of the lipid in a real cell. Regardless, these data suggest that head groups more likely to harbor a negative charge are more effective at potentiating ASIC3.

The two bulkier head groups showed differing effects. Ethanolamide phosphate, which does have a site likely to have a negative charge over the range of pH used here, shifted the pH_0.5_ to a similar degree as the carboxyl head, while the taurine head group, despite its low pKa, had no effect on channel activation (Fig. 2B). This reduced efficacy may stem from steric clashes between these bulkier head groups and the binding site on the channel. Taken together, these data indicate that the charge of the head group can have a profound impact on the ability of the lipid to potentiate ASIC3 currents.

### A longer acyl tail is important for PUFA potentiation of ASIC3

Next, we sought to determine how the structure of the tail impacts regulation of ASICs. PUFA tails are often broken down into four properties: length, number of double bonds, position of the double bonds, and ω-number. To understand how each of these contribute to PUFA modulation of ASICs, we examined a total of 24 PUFAs with identical carboxyl head groups (Table 1). We again measured changes in the pH dependence of activation in response to 10μM PUFA. Figure 3A shows PUFAs with varying tail properties and their respective shifts in the activation pH_0.5_ which ranged from a 0.32 pH unit alkaline shift all the way to a 0.07 acidic shift. These data indicate that the nature of the tail can have a considerable impact on the modulation of the channel.

**Figure. 3.**
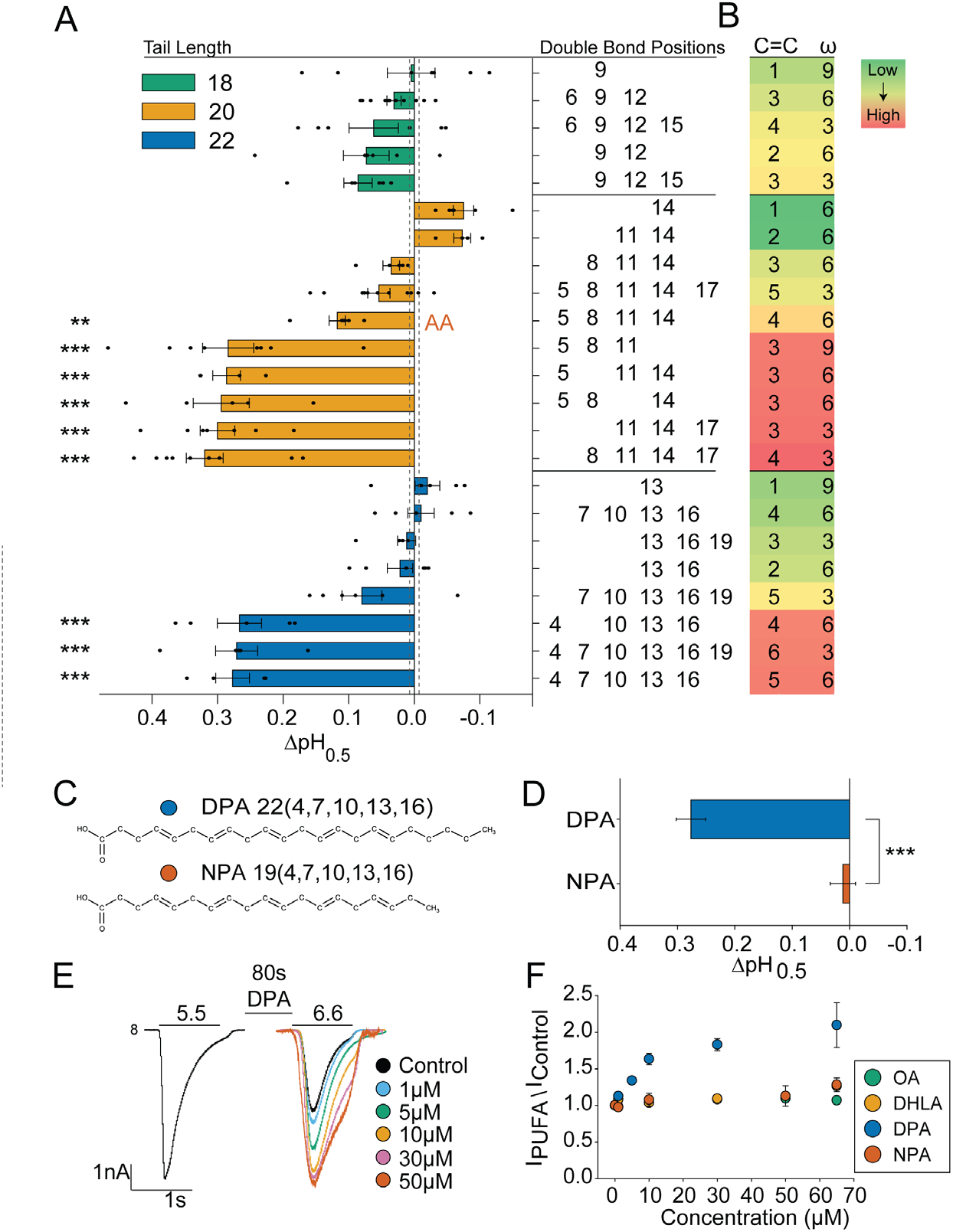
Effects of PUFA tail properties on ASIC3 potentiation. (A) Activation ΔpH_0.5_ values for ASIC3 induced by 10μM of PUFAs with the indicated tails. Tails are grouped based on their tail length and then ordered based on the magnitude of their ΔpH_0.5_ shifts. The exact positions of the tail double bonds are given on the y-axis. (n = 4-11, Data in Table 1). Vertical dashed lines indicate SEM of control (n = 70). (B) Heat map showing various tail properties (length, number of double bonds, and ω-number) for the data in panel A. Each row in panel B corresponds to the PUFA adjacent in panel A. Dark red indicates the largest alkaline ΔpH_0.5_ measured while dark green indicates a small acidic shift. (C) Structures and (D) ΔpH_0.5_ values for DPA and NPA which have identical double bond positions but differing acyl tail lengths (pH_0.5_ = 6.83 ± 0.03, n = 5 for DPA; and 6.57 ± 0.02, n = 6 for NPA). (E) Representative traces for a single cell showing the protocol used to generate data in F. Current induced by 1s acidic jumps to pH 5.5 and 6.6 was measured for the control followed by the measurement of current at pH 6.6 after 80s incubation of increasing concentrations of DPA at a resting pH of 8. (F) Plot showing the fold-increase in current measured at pH 6.6 plotted as a function of concentration for OA, DHLA, DPA, and NPA (n = 4-11). All data given as mean ± SEM. Asterisks in A represent p-values as determined by post-hoc Dunnett’s test. Asterisks in D represent p-values as determined by unpaired t-test. * p < 0.05, ** p < 0.01, *** p < 0.001 (see methods for details).

In determining the tail properties important in PUFA potentiation, we first examined tail length. All PUFAs with an 18-carbon acyl tail had small (< 0.1 pH unit), non-significant, effects, whereas many of the 20- and 22-carbon tail PUFAs shifted ASIC3 activation pH_0.5_ to a greater extent with several shifting by more than 0.26 pH units (Fig. 3A). This can also be seen in the heat map in figure 3B. Each column of the map represents an individual tail characteristic of the adjacent PUFA in figure 3A. The magnitude of the shift in activation at 10μM is illustrated by the color scale ranging from dark red (strong alkaline shifts) to dark green (small acidic shift). Although a longer tail length appeared necessary to confer stronger regulation, it was not sufficient. Several PUFAs with 20- and 22-length acyl tails failed to potentiate ASIC3, indicating other tail properties are important to PUFA potentiation.

### PUFAs with a longer acyl tail require more double bonds to potentiate ASIC3

Our data suggest that a minimum of three double bonds in the PUFA tail were necessary for strong potentiation (Fig. 3A, 3B). No PUFAs with either one or two double bonds potentiated ASIC3 currents. Each tail length appears to have an optimal window of required double bonds to effectively potentiate ASIC3. We observed that as the acyl tail got longer, the range of optimal double bonds also increased. For PUFAs with a 20-length tail, strong potentiators all had three to four double bonds. For PUFAs with 22-carbon tails, four to six double bonds were necessary for strong potentiation of ASIC3 (Fig. 3A, 3B).

To look at this more carefully, we measured the shift in activation pH_0.5_ caused by docosapentaenoic acid (DPA) and nonadecapentaenoic acid (NPA) which share the identical number and position of double bonds but differ in tail length ([22(4,7,10,13,16)] and [19(4,7,10,13,16)], respectively) (Fig. 3C). At 10μM, the 22-carbon tail DPA strongly potentiated ASIC3 (ΔpH_0.5_ = 0.27), whereas the 19-carbon NPA, had no effect on the channel (ΔpH_0.5_ = 0.01) (Fig. 3D). These data are consistent with our observations. First, tails of at least 20 carbons are necessary, but not sufficient for strong potentiation. Second, having five double bonds in a longer acyl (22-carbon) tail is compatible with strong modulation, but having the same five double bonds does not result in a good potentiator for shorter tails.

To ensure the differences observed for NPA and other PUFAs that did not alter the activation pH_0.5_ was not simply from an increase in EC_50_ values, we compared the concentration dependent effects for both weak and strong ASIC3 potentiators. To do this we measured ASIC3 currents elicited by brief (1s) pulses to the ~pH_0.5_ (6.6) and applied increasing concentrations of PUFA to the same cell (Fig. 3E). The black trace in figure 3E shows fully activated ASIC3 current at pH 5.5. The overlayed traces to the right show currents elicited with a pulse from pH 8 to pH 6.6 at increasing concentrations of DPA. These data show that with increasing concentrations of DPA the channel can be maximally activated even at pH 6.6. In contrast, NPA, oleic acid (OA) [18(9)] and dihomo-γ-linolenic acid (DHLA) [20(8,11,14)] which all failed to significantly shift the activation pH in figure 3A also showed no further potentiation of pH 6.6 currents at higher concentrations (Fig. 3F).

### A double bond near the head group is important for 22-length PUFAs

For PUFAs with 18- or 20-length tails, there was no specific double bond position that appeared necessary for potentiation. However, for the 22-length PUFAs, a double bond at position four was present in every strongly potentiating PUFA (Fig. 3A, 3B). DPA [22(7,10,13,16,19)] did not significantly affect ASIC3 while DPA [22(4,7,10,13,16)] with a double bond at position four, had a large (ΔpH_0.5_ = 0.27 pH units) alkaline shift in the activation pH, despite sharing four out of five double bond positions. Likewise, docosatetraenoic acid [22(7,10,13,16)] failed to alter ASIC3 currents despite having all the same double bonds as DPA [22(4,7,10,13,16)] except for the four-position.

Finally, we examined the relationship between ω-number and the functional effect of the lipid. The ω-number refers to the number of carbons away from the methyl end of the tail that the last double bond appears. Previous work has shown that the ω-number correlates with some effects that PUFAs have on other classes of ion channels^19^. However, there was no relationship between ω-number and the shift in the pH dependence of activation of ASIC3 by PUFAs (Fig. 3B).

### Amino acid head groups have larger magnitude effects on ASIC3 currents

Our data to this point has shown that the structure of the head and tail groups impact the magnitude of the effect that the lipid has on ASIC3 function. In principle, this impact can stem from a change in the binding or a change in the ability of the lipid to alter ASIC3 function. Thus, we set out to examine how the different structural features of the lipids impacted the magnitude of the effect as well as the binding to ASIC3.

First, we compared the EC_50_ of the shift in the pH dependence of activation for AA, DHA, and AG (Fig. 4A). AG and DHA showed a modestly lower EC_50_ values of 5.40μM ± 1.12μM and 4.9μM ± 0.66μM, respectively, compared to 11.62μM ± 2.08μM for AA. However, DHA and AA, both with carboxyl head groups had comparable maximum shifts while AG had a larger maximum shift in the pH_0.5_ (Fig. 4A). Again, while all three of these lipids caused strong shifts in the activation pH, there was little shift in the pH dependence of desensitization (Fig. 4B).

**Figure. 4.**
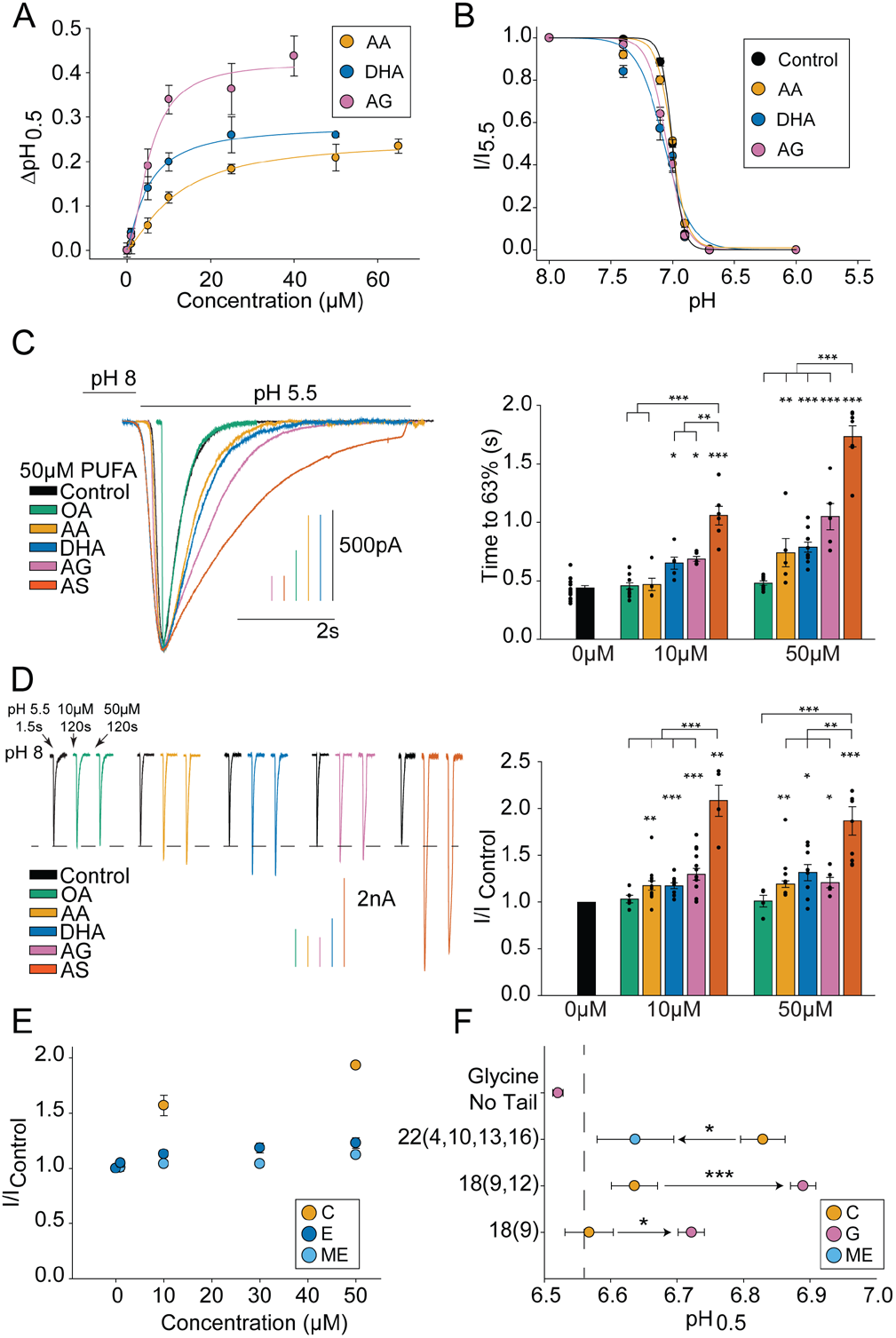
N-acyl amino acids with the arachidonic tail more strongly potentiate ASIC3 currents compared to AA. (A) Activation ΔpH_0.5_ values plotted as a function of concentration for AA, DHA, and AG (EC_50_ = 11μM ± 2μM, n = 3-7 for AA; 5μM ± 1μM, n = 4-7 for DHA; and 5μM ± 1μM, n = 3-6 for AG.) AA replotted from Fig. 1 for comparison. (B) Plot showing the pH dependence of desensitization in the presence of 10μM AA, DHA, and AG (pH_0.5_ = 7.07 ± 0.04, hill slope = 4 ± 1.6, n = 6 for DHA; and pH_0.5_ = 7.05 ± 0.01, hill slope = 5.7 ± 0.96, n = 5 for AG). Control and AA replotted from Fig. 1 for comparison. (C) Representative traces showing the desensitization of ASIC3 currents evoked by an acidic jump to pH 5.5 in the presence of 50μM of various lipids (left). Plot showing ASIC3 desensitization rates for different lipids at 10μM and 50μM. Rates are given as the time to reach 63% decay of the peak of current measured at pH 5.5 (n = 5-19) (right). (D) Representative traces showing the change in ASIC3’s current magnitude measured at pH 5.5 in the presence of various lipids (left). Each group of three traces represents the current measured for a single cell prior to lipid application followed by subsequent 2min exposures to 10μM and then 50μM lipid. Plot showing the fractional increase in ASIC3 current measured at pH 5.5 for various lipids at 10μM and 50μM (n = 5-13) (right). (E) Plot showing the fold-increase in current measured at pH 6.6 plotted as a function of lipid concentration for AA and two AA-derivatives with neutral head groups, E and ME (n = 4-7). (F) Plot showing the activation pH_0.5_ values of ASIC3 measured in the presence 10μM of the indicated PUFA with native carboxyl head and their corresponding head group derivatives. Glycine molecule without an acyl tail is also shown measured at 100μM glycine. Tail length and double bond position is indicated on the y-axis and head group is indicated in the color key (C = carboxyl, G = glycine, ME = methyl ester). Arrows denote the shift from the carboxyl head group to the substituted head group. Vertical dashed line represents the activation pH_0.5_ of ASIC3 Control. All data given as mean ± SEM. Asterisks above bars in D and F represent p-values as determined by post-hoc Dunnett’s test. Asterisks for between group comparisons in D and F as determined by post-hoc Tukey’s test. Asterisks in H represent p-values as determined by unpaired t-tests. * p < 0.05, ** p < 0.01, *** p < 0.001 (see methods for details).

In addition, the more negatively charged head groups also had a larger impact on the desensitization rate of ASIC3 (Fig. 4C). All potentiating lipids tested also slowed the rate of desensitization in a concentration dependent manner (Fig. 4C). OA, which had no effect on the pH_0.5_, also had no effect on the rate of desensitization. Again, the NAAAs had the largest impact on the rate of desensitization at either 10μM or 50μM. In fact, AS was significantly stronger at slowing desensitization compared to all other lipids tested, including AG.

A more complicated result was seen looking at the changes in I_MAX_ that these lipids elicited. These currents were measured by switching between pH 8 and pH 5.5 for 1.5s and then repeating this measure after 120s exposure to the lipid (Fig. 4D). Again, at both 10μM and 50μM, application of OA had no effect on I_MAX_, while all potentiating lipids significantly increased the current at pH 5.5. Surprisingly, unlike the observed effects on the rate of desensitization and the pH_0.5_, the increase in I_MAX_ was the same for all potentiating lipids (~1.2-fold) apart from AS, which was significantly greater than all other lipids, producing an ~2-fold increase in I_MAX_. (Fig. 4D). Also interesting was that each lipids effect on I_MAX_ was not significantly different at 10μM and 50μM whereas both the slowing of desensitization and shift in activation pH_0.5_ increased going from 10μM to 50μM.

While the NAAAs were stronger potentiators, lipids with neutral head groups had minimal effects on ASIC3 currents even at high concentrations. To show this, we compared the concentration dependent effects of AA to two weakly potentiating AA-derivatives, methyl ester and ethanolamide, following the same protocol illustrated in figure 3E where we pulse to a single pH (6.6) and add increasing concentrations of PUFA. AA shows a clear concentration dependent increase in pH 6.6 evoked ASIC3 currents while the neutral ethanolamide and methyl ester head groups produce lipids that are weak potentiators of ASIC3 currents even at increasing concentrations of lipid (Fig. 4E).

Given this ability of the head group to substantially impact the magnitude of the effect the lipid has on ASIC3 currents, we wondered whether we could convert a poor potentiator into a strong one by only changing the head group to one with a lower pKa, or conversely reduce the effect of a strong potentiator by replacing the head group with a neutral one. To do this we first compared docosatetraenoic acid [22(4,10,13,16)] with carboxyl head group to a methyl ester head group derivative with the same tail and found that the potentiating effect was significantly reduced with the methyl ester (Fig. 4H). Conversely, both linoleic acid [18(9,12)] and oleic acid [18(9)], which had no statistically significant effect on ASIC3 currents, could both be converted into stronger potentiators upon replacement of their carboxyl head groups with a glycine (Fig. 4F). Although the glycine head group enhanced potentiation of all PUFAs tested, addition of glycine alone (no tail) showed no potentiation, even up to 100μM, illustrating that an acyl tail is required (Fig. 4F). These data show that the head group can convert a lipid from a bad potentiator to a good one or vice versa suggesting that the head group is likely the critical determinant of the strength of potentiation.

### Arachidonoyl glycine can act as a ligand for ASIC3 even at neutral pH

In addition to the effects on ASIC3 currents we have reported so far, AG created a non-desensitizing ASIC3 current at modest acidifications. Figure 5A shows representative currents elicited from 20s pulses to pH 7.1 from cells in control conditions and incubated with 10μM AG. The presence of AG results in a current that is ~1.4% of the peak current and shows no signs of desensitizing even at 20s. This current likely arises from a “window current” mechanism. The term window current refers to a range of pH values where the channel may activate but not desensitize. The alkaline shift in the activation pH_0.5_ without a concomitant impact on the desensitization pH_0.5_ opens a window of pH over which we would predict a non-desensitizing current might arise from application of lipids like AG. This window current idea predicts that the magnitude of the non-desensitizing current should show a biphasic dependence on pH increasing and then decreasing with progressive acidification. This is exactly what we see for DHA and AG. Figure 5B shows the magnitude of the non-desensitizing current as a function of pH. Both lipids increase the size of the non-desensitizing current with the stronger potentiator, AG, having a larger effect.

**Figure. 5.**
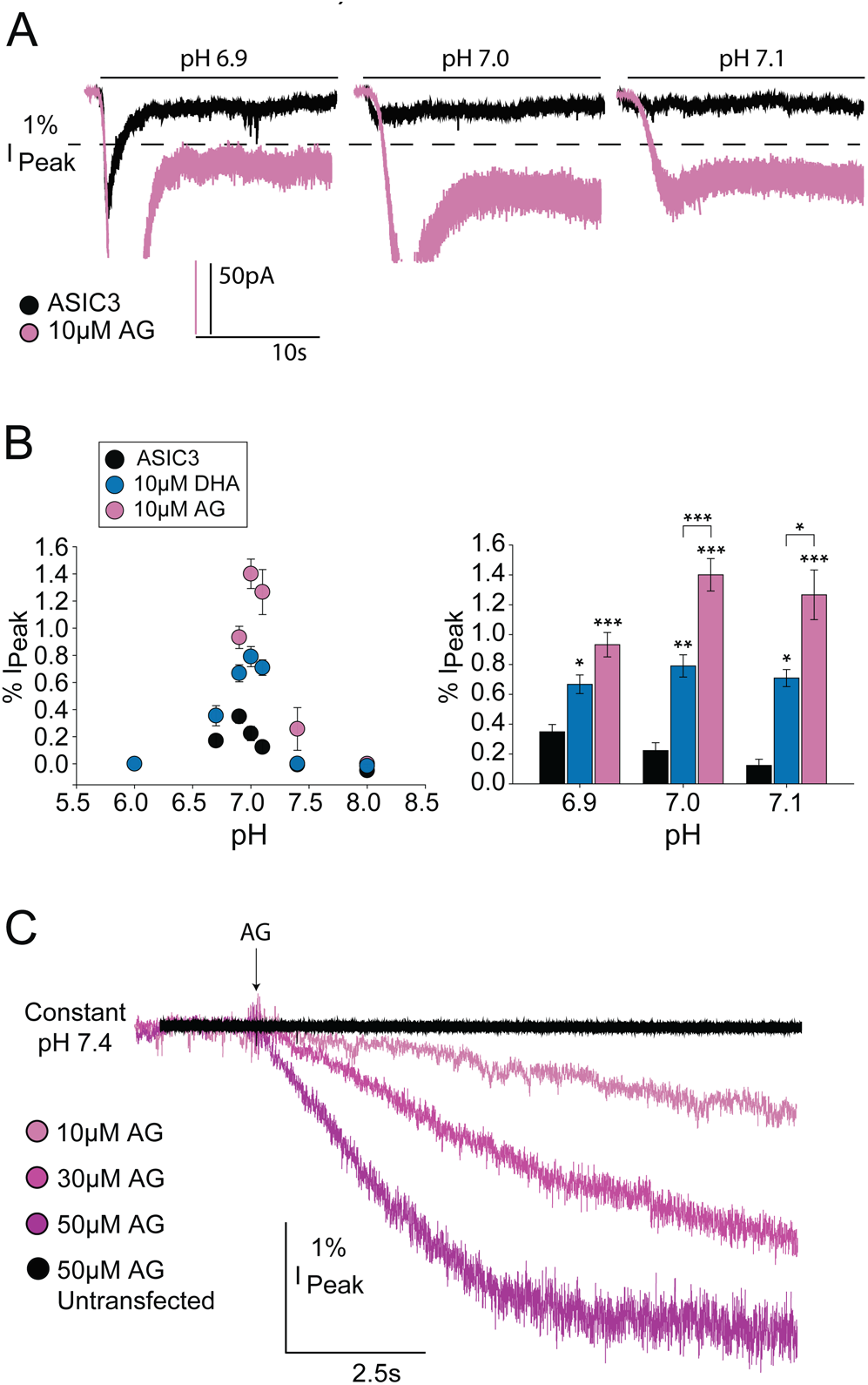
Potentiating lipids cause a non-desensitizing current during moderate acidosis. (A) Representative currents evoked by 20s of indicated for pH for ASIC3 ±10μM AG. The dotted line represents a current value that is 1% of the peak (pH 5.5) current. (B) (left) Plot showing the sustained current generated as a function of pH for control (n = 9), DHA (n = 4), and AG (n = 6) and (right) the same data replotted for pH 6.9, 7.0 and 7.1 to better show comparisons of effects. Sustained current is measured as the 5s average of the current 15s after pH application and is shown as the percentage of the peak current measured at pH 5.5. (C) Representative ASIC3 currents evoked by application of different concentrations of AG while maintaining a constant pH of 7.4. Trace shown in black represents the application of 50μM AG to untransfected cells under identical conditions. All data given as mean ± SEM. Asterisks in B as determined by post-hoc Tukey’s test with the above bar asterisks representing significance as compared to ASIC3 control. * p < 0.05, ** p < 0.01, *** p < 0.001 (see methods for details).

This mechanism may allow these stronger lipids to act as a ligand even at neutral pH. To look at this, we held our cells expressing ASIC3 at pH 7.4 with constant perfusion of control solution and made a rapid switch to solution containing various concentrations of AG. This resulted in a lipid evoked current in ASIC3 that increased with increasing concentration of AG (Fig. 5C). Application of AG to cells not expressing ASIC3 showed no current (Fig. 5C, black).

### Head group structure impacts wash on/off of the lipid

Despite having nearly identical EC_50_ values, the wash on and off kinetics of DHA and AG differed substantially. To test the wash on rate, we pulse to pH 6.6 for 1s with 3s (for AG) or 5s (for all other lipids) returns to pH 8 to allow for recovery from desensitization. After three pulses in control solutions, the cell is then continuously exposed to lipid for the duration of the wash on experiment (Fig. 6A). Solutions are then switched back to control (no lipid) and wash off is measured using the same protocol. The time course of wash on, as determined by the amount of time to reach 1/e of the maximum effect (see methods), for AG was 5.97s ± 0.73s, approximately six times faster than either AA or DHA, 31.08s ± 4.96s and 29.20s ± 4.98s, respectively (Fig.6B,D). The time to reach 63% wash off for AG, 15.35s ± 0.82s, was two times faster than AA or DHA, 30.18s ± 3.00s and 30.53s ± 1.01s, respectively (Fig. 6C,D). However, unlike AA and DHA, AG showed complete wash off while DHA and AA plateaued at values approximately 20% greater than control pH 6.6 values. While none of these results are direct measures of a change in affinity, they are consistent with the idea that the structure of the lipid may alter the binding to the channel which is an expected result.

**Figure. 6.**
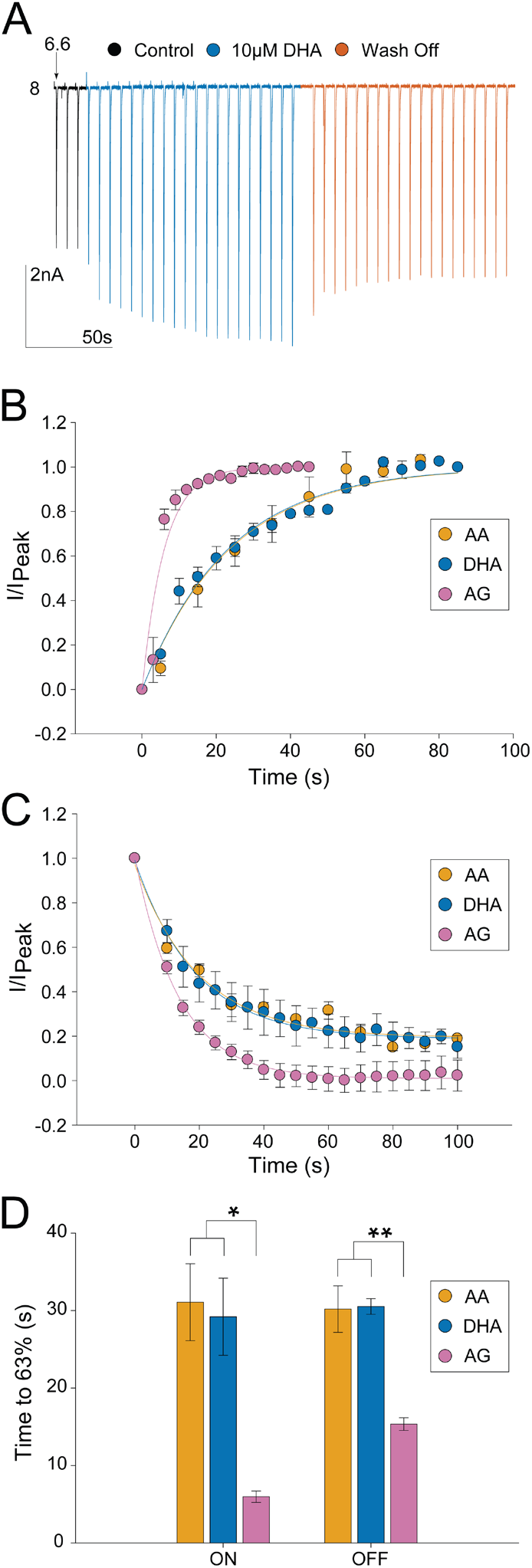
Time course of lipid wash on/off of ASIC3. (A) Representative traces for a single cell showing repeated pH 6.6-evoked currents upon addition and removal of 10μM DHA. (B) Wash on and (C) wash off time courses for AA, DHA, and AG. Current was normalized by first subtracting the control current from each time point and then dividing by the maximal current response measured at pH 6.6 at the end of wash on. (C) Currents at time 0s in C represents maximal current reached for wash on in B. Currents measured during AA and DHA wash off were not able to return to control levels (prior to wash on). (D) Bar plot showing time to reach 63% maximal current response (ON) or time to diminish maximal current response by 63% (OFF) as determined by fits (n = 3-5) (see methods). All data given as mean ± SEM. Asterisks in D represent p-values for between group comparisons as determined by post-hoc Tukey’s test. * p < 0.05, ** p < 0.01, *** p < 0.001 (see methods for details).

### An arginine at the top of TM1 of ASICs is critical for PUFA potentiation

Given that the charge of the headgroup impacted the strength of lipid potentiation of ASIC3, we hypothesized that there may be residues on the channel near the membrane boundary that are able to make interactions with the charged head groups of the lipid. This has been demonstrated for other channels where interactions occur with Y, R, or K residues^12,16,20,33,35^. We therefore sought to test whether PUFAs modulate ASICs through similar interactions.

We attempted to narrow down the region on the protein important for regulation before making more targeted mutations. First, we examined the sidedness of the PUFA regulation. To do this, we added 10μM AG to the patch pipette and saw no change in current over one minute nor was there any change in the rate of desensitization as compared to control values (Fig. 7A, 7B). Addition of 10μM AG to the extracellular side of the same cell resulted in a rapid increase in both the current as well as a slowing of desensitization (Fig. 7A, 7B). In addition, inclusion of 10μM AG in the patch pipette did not affect the activation pH_0.5_, even after 10 minutes of recording despite steady state being reached within 10s for external application (Fig. 7C, for wash on rates see Fig 6D and 7A). We next showed that 10μM DHA resulted in an alkaline shift in the pH dependence of activation of ASIC1a (Fig. 7D). Together these data suggest that PUFAs act on the extracellular leaflet of the plasma membrane and likely require residues conserved between ASIC1a and ASIC3. We found five candidate residues as possible interaction sites (Fig. 7E, 7F). We made 11 channel mutants that would neutralize the potential interaction with the side chain. Not all the mutations produced functional channels (Supplementary Table 1).

**Figure. 7.**
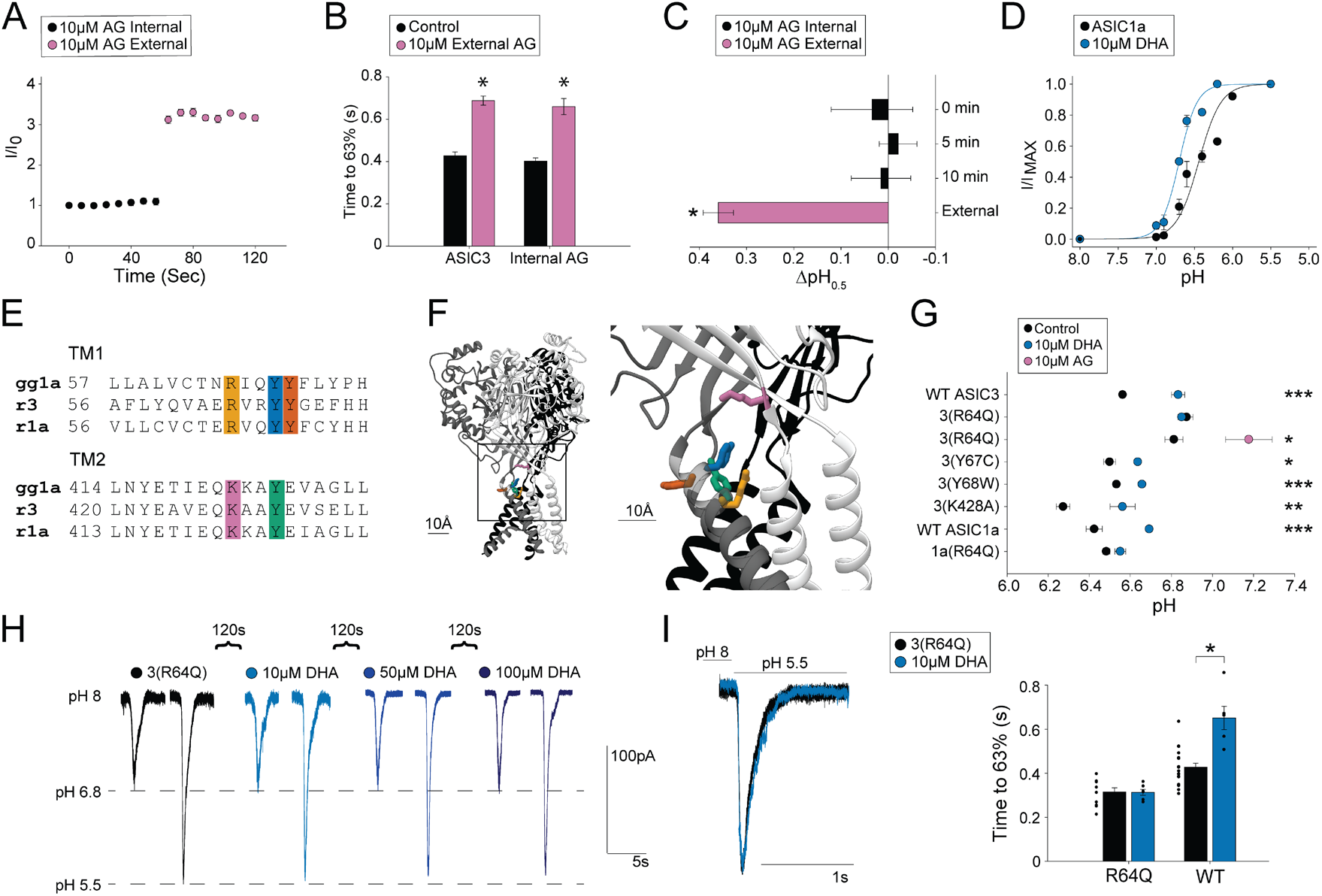
PUFA potentiation of ASICs is dependent on an arginine in the outer segment of TM1. (A) Time course of ASIC3 activations by repeated exposures to pH 6.6 from pH 8. Internal application represents 10μM AG within patch pipette, this was followed by activations in the presence of 10μM AG in the external solutions (n = 3). (B) Plot showing desensitization rates for ASIC3 ± 10μM internal AG before and after application of 10μM external AG (n = 3). ASIC3 measurements in the absence of internal AG are replotted from Fig. 4 for comparison. Rates are given as the time to reach 63% decay of the peak of current measured at pH 5.5. (C) ΔpH_0.5_ values of ASIC3 activation over time with 10μM AG within the patch pipette. 0 min denotes the pH_0.5_ measured immediately after going whole-cell compared to control pH_0.5_ values (n = 3). (D) pH dependence of activation for ASIC1a ± 10μM DHA (pH_0.5_ = 6.43 ± 0.05, hill slope = 2.3 ± 0.49, n = 7 for Control; and pH_0.5_ = 6.70 ± 0.02, hill slope = 3.7 ± 0.88, n = 6 for DHA). (E) Sequence alignment for the extracellular segments of TM1 and TM2 for chicken ASIC1, rat ASIC1a, and rat ASIC3. Highlighted residues represent potential PUFA regulation sites. (F) Structure of chicken ASIC1 showing highlighted residues from E. Structure visualized using Chimera 1.12 and PDB 6VTK. (G) Activation pH_0.5_ values for ASIC mutants ± 10μM DHA or 10μM AG (n = 4-10, Data given in Supplementary Table 1). WT ASIC3 values replotted from figure 3 for comparison. (H) Representative traces for a single cell showing currents evoked by pH 6.8 and pH 5.5 for ASIC3 with R64Q mutation before and after addition of increasing concentrations of DHA. (I) (left) Representative traces showing pH 5.5 evoked currents from ASIC3 with R64Q mutation ± 10μM DHA and (right) plot showing ASIC3 WT and ASIC3 with R64Q mutation desensitization rates ± 10μM DHA (n = 6-10). Rates are given as the time to reach 63% decay of the peak of current measured at pH 5.5. WT rates replotted from Fig. 4 for comparison. All data given as mean ± SEM. Asterisks in B for internal AG represent p-value as determined by paired t-test. Asterisks in C represent the p-value as determined by post-hoc Dunnett’s test. Asterisks in G represent the p-value as determined by unpaired t-tests between Control and DHA measurement for each construct. Asterisks in I represent the p-value as determined from post-hoc Dunnett’s test in Fig. 4 for DHA and the p-value determined by an unpaired t-test between R64Q ± Lipid. * p < 0.05, ** p < 0.01, *** p < 0.001 (see methods for details).

We tested the effect of DHA on these mutants due to its stronger potentiation of ASIC3, rationalizing that we would more easily be able to see reductions in PUFA effects. We found functional mutations in four of the five positions and measured whether 10μM DHA altered the pH dependence of activation in these mutants. A single position, R64Q, eliminated the shift in the activation pH_0.5_ produced by DHA (Fig. 7G). For the Y67C and Y68W mutants, DHA still potentiated the currents, but the effect was reduced compared to WT potentiation. Looking at the structure of ASIC1 from chicken, R64, Y67, and Y431 (R65, Y68, and Y425 in the chicken), are all within 10Å of one another (Fig. 7F). Mutation of the more distal K428, which resides in the β12 strand, to an alanine yielded functional channels that were still fully potentiated by DHA (Fig. 7G). To further confirm that this region of the channel is important for PUFA potentiation of ASICs, we also mutated R64 in ASIC1a. Like with ASIC3, 10μM DHA did not significantly affect the pH dependence of activation of ASIC1a with the R64Q mutation (Fig. 7G). To further confirm the importance of R64, we looked at the ability of higher concentrations of DHA to potentiate ASIC3 R64Q currents. To do this, we again held our cells at pH 8 and pulsed to pH 6.8 (~pH_0.5_ of the mutant channel) and then to pH 5.5. Even at 100μM DHA we saw no effect on the current at pH 6.8 or pH 5.5 suggesting that the mutation eliminated both the shift in pH_0.5_ and the change in I_MAX_ measured in WT channels (Fig. 7H). In addition, the ASIC3(R64Q) mutation also eliminated DHA’s slowing of desensitization (Fig. 7I). Interestingly, AG was still able to shift the pH dependence of R64Q channels to the same extent as WT channels (Fig. 7G, pink circle). This suggests the possibility that R64 is part of the binding site but interacts with the head group in a part of the head group where the structures differ.

## Discussion

The results of our study provide the first mechanistic insight into lipid regulation of ASICs. Previous studies have shown that AA can potentiate ASIC currents via a shift in the pH dependence of activation as well sensitize neurons to pH dependent action potential firing^25–28^. However, this work is the first to show that this potentiation can result from the binding of a number of different lipids beyond AA. In fact, several endogenous PUFAs and lipids have a stronger potentiating effect on the channel. In addition, we show that these lipids can act to stabilize the open state of the channel which manifests as a slowing of the rate of desensitization and an increase in the current even at saturating pH.

A previous study examined AA along with four other related lipids and concluded that the head group had minimal impact on potentiation and that increasing the number of double bonds leads to an increased effect on ASICs^26^. Here we tested 24 PUFAs with the carboxyl head group and differing tails. Our data do not support the hypothesis that increasing numbers of double bonds are the main determinant of potentiation. First, we found that shorter 18-length tails showed weak to no potentiation and longer 20- or 22-length tails were necessary, but not sufficient, to confer strong potentiation. Our data suggests that longer tails required more double bonds to potentiate ASIC3 currents but that too many double bonds could also reduce efficacy (see 20[5,8,11,14,17] in Fig. 3). For 22-length PUFAs, a double bond at position four also seemed to be important for strong potentiation of the channel. The double bonds change the shape and flexibility of the tail and so it is easy to imagine that the more effective PUFAs have tails whose structure more effectively sit in its binding site on the channel, but more work is needed to understand how the precise structure of the tail impacts PUFA action^36,37^.

Again, in contrast to previous findings, we show here that head groups with lower pKa such as the NAAAs arachidonoyl glycine and arachidonoyl serine had the largest effects ASIC3^26^. Perhaps most strikingly, we were able to convert weakly potentiating PUFAs like linoleic acid [18(9,12)] and oleic acid [18(9)] into significantly stronger potentiators by replacing the carboxyl head group for a glycine. Conversely, we could significantly decrease the ability of docosatetraenoic acid [22(4,10,13,16)] to potentiate ASIC3 current by substituting a methyl ester head group for the normal carboxyl. We believe that these data suggest that the head group of these lipids is the major determinant of the strength of the lipid modulation of the channel and that the structure of the head group can toggle the strength of the potentiation across a multitude of tail structures.

Consistent with the idea that the head group is critical for modulating ASIC currents, mutating R64 eliminated DHA-dependent potentiation of ASIC3 currents. Similarly, mutation of neighboring residues showed a reduction of DHA potentiation, while a more distal mutation at K428 had no effect. This suggests the possibility that these residues around R64 create a binding surface for the lipid head group on the channel. It is difficult to distinguish between the possibility that R64 is important for binding versus important for coupling the binding of lipid to the gate of the channel. However, we believe that several pieces of evidence argue for R64 being important for PUFA binding to the channel.

First, we were able to show that these lipids act on the outer leaflet of the plasma membrane where R64 resides. Second, R64 is conserved in ASIC1a which is also potentiated by PUFAs. Third, DHA failed to act on R64Q even at concentrations as high as 100μM DHA. Finally, mutation of R64 eliminated the effect of DHA but did not impact the potentiation by AG. Assuming these two lipids occupy approximately the same binding site and act via a conserved mechanism, if R64 was involved in the coupling of the lipid to channel opening you would predict a comparable effect of the mutation on the potentiation by each lipid. Since the mutation has a strikingly different effect on the two lipids it is likely that R64 is in the binding site and interacts with the head group in a position where the head group structure differs between the lipids. This argument has been made previously for understanding the binding site for cGMP on CNG channels^38^.

Interestingly, two recent structures of ASIC1a from both chicken and human shows endogenous lipid present between TM1 and TM2 of the same subunit immediately adjacent to our putative binding site^21,39^. Complicating our hypothesis, R64 points into the pore of the channel in the solved structures^22–24^. However, MD simulation of this site suggest that the R64 position undergoes significant rearrangements during gating and there is considerable evidence that lipid head groups can interact with residues even on the inside of the pore^40–42^.

Furthermore, there is precedent in other channels for this sort of interaction between a negatively charged head group and a positively charged side chain. K_v_7.1 regulation by PUFAs is strengthened by head groups with lower pKa^14,16,18,20^. Recent work has suggested that the negatively charged head group interacts with an arginine and a lysine residue on the extracellular side of the voltage sensor and pore domain of K_v_7.1^43^. In addition, DHA binds to and speeds the desensitization of the pH gated GLIC channel^33^. A crystal structure of the channel in the presence of DHA solved at pH 4, showed the carboxyl head group forming a salt bridge with the guanidinium group of an arginine residue near the extracellular side of a transmembrane helix^33^.

However, we cannot conclusively say that R64 is in the binding site for the lipid head group on ASICs. An alternative hypothesis would be needed if AG and PUFAs acted via different structural mechanisms. Future work using molecular dynamics or solved structures in the presence of these lipid will be needed to confirm our hypothesis.

How binding to this putative interaction site alters channel function will be important to examine in future studies. There are a number of potential mechanisms. First, mutation of this arginine alters gating of ASIC1a suggesting that the lipids could simply be altering the conformational dynamics of this regions^41^. Second, work in K_V_1.4 channels has suggested that PUFA head groups can change the pKa of nearby protonatable residues by altering the electrostatic potential near the lipid binding site^44^. Two nearby histidine residues have been suggested as candidate sites for pH sensing in ASIC1a and their ability to be protonated may be altered by lipid binding^45^. Finally, Ca^2+^ binding to ASIC3 is thought to inhibit the channel and there are several studies that propose the idea that the proton dependence of the channel stems from protons competing off Ca^2+^ ions to unblock the channel as opposed to protons causing a conformational change that opens a gate^46,47^. R64Q is located very close to these putative Ca^2+^ binding residues, E425 and D432, and may even form a salt bridge with E425^22,48^. It is possible that the lipids we examined here may be altering the putative Ca^2+^ binding site and thus the gating of the channel.

Perhaps most relevant to the physiological function of ASICs, the strongly potentiating NAAAs, like AG, were able to act as a ligand for the channel at neutral pH activating a small but non-desensitizing ASIC3 current. The ability of these NAAAs to act as a ligand likely owes to their ability to create a “window current”. This window current mechanism also means that there can be significant non-desensitizing ASIC currents at modest acidifications (pH 7.1) as well. It is still a matter of debate what pH ASICs are likely to experience at synapses. Work in retinal bipolar cells has shown that the synapse may acidify by 0.2-0.6 pH units^49^. In our experiments, a pH of ~6.6 is required to activate half of the ASIC3 current and a pH of 6.9 would activate almost no ASIC3 currents. However, in the presence of some of the lipids studied here, modest acidification would result in small but non-desensitizing ASIC3 currents suggesting that lipids may be able to activate a small persistent ASIC current in neurons. These persistent currents could have significant impacts on neuronal excitability. A previous study showed that AA increased the pH dependent excitability of skin DRG neurons^26^. Our data would predict that lipids like DHA, AG, and AS would have potentially even larger effects on neuronal excitability.

The physiologically relevant concentration of these lipids in the extracellular space or plasma membrane of cells is not well known, however, concentration of unesterified PUFA in plasma has been suggested to be in the 10μM to 50μM range^50–52^. This concentration can increase to as high as 400μM depending on diet and can also increase in many pathological conditions including inflammation, ischemia, and epilepsy^50,52–54^. Reports have shown that DHA may represent 8-18% of the fatty acids in the adult brain and make up an astounding 50-70% of the fatty acids found in the rod outer segment in the retina^55,56^. NAAAs like AG have been demonstrated to be important bioactive lipids that regulate ion channels^57^.

In summary, we have shown that PUFAs and many of their derivatives can increase ASIC3 currents by shifting the pH dependence of activation and stabilizing the open state of the channel. Lipids with head groups with a low pKa, like the NAAAs, are potent activators of ASIC3. These head groups can convert a poorly potentiating lipid into a strongly potentiating one. In addition, the NAAAs can activate the channel even at neutral pH. Our data are consistent with a model where the acyl tail interacts with the core of the TM region of ASIC3 while the head group interacts in a binding site near the outer portion of the TM segments. While more work is needed to confirm this idea, we hypothesize that an arginine (R64) near the outer leaflet of TM1 is in the binding site for the head group and interacts with the carboxyl head group of PUFAs but fails to interact with the NAAA head groups. This idea impacts the mechanism of lipid regulation of ASICs more broadly. Lysophophatidyl choline (LPC) was shown to activate ASICs at neutral pH as well^28^. The lipids are structurally quite distinct as the effective LPCs are LPC(16:0) and LPC(18:1) which have short acyl tails and either zero or one double bond. However, the head group is zwitterionic and contains a negatively charged phosphate group. It will be interesting to find out if these lipids act through the same binding site or if there are multiple potential sites for modulating channel function. In addition, PUFAs are potential lead compounds for the modulation of ASIC function. Our work identified two potential inhibitors, eicosenoic acid [20(14)] and eicosadienoic acid [20(11,14)] that will require future investigation. We have thus identified the basic principles for modulation of ASIC3 by PUFAs and PUFA derivatives which can form the basis for the design of new compounds to alter ASIC function in neurons by targeting the site where this modulation occurs.

## Materials and Methods

### PUFA Nomenclature

For common PUFAs (AA, OA, DHA, etc.) abbreviations are used after being defined in the text. All abbreviated PUFAs can be found in Supplementary Table 2. Other PUFAs are named using their tail properties in the format of [length(position of double bonds)]. All acyl tail doubles bonds are in the cis configuration for PUFAs and PUFA-derivatives used in this study. Unless otherwise specified, all PUFAs used in this study contain a carboxyl head group. A full list of the PUFAs used in this study with their head groups, tail lengths and position of doubles bonds is given in Table 1.

### Materials and Mutagenesis

PUFAs used in this study were purchased through Cayman Chemical. Plasmids for ASIC1a, and ASIC3 each from rat were gifts from David Julius and subcloned into pcDNA3.1. To visualize cell expression, WT, and mutant ASICs a short proline rich linker was used to join mCerulean3 to the C-terminus of the channel, previously reported to have minimal effects on channel gating^58^. The mutants used in this study were made using site-directed mutagenesis either in house or through Biozilla services.

### Cell Culture and Transfection

Chinese hamster ovary (CHO-K1) cells (ATCC) were cultured in Ham’s F12 media with 10% FBS at 37 °C in 5% CO2. Cells at ~70% confluency were transfected with indicated ASIC plasmid DNA. Transfection was achieved either via electroporation with a Lonza 4D Nucleofector unit or through chemical means using Mirus TransIT-Transfection Reagent (Mirus) following the manufacturer’s protocols. Following transfection, cells were plated on 12 mm glass coverslips coated in poly-L-lysine.

### Electrophysiological Recordings

All experiments were performed in the whole-cell patch-clamp configuration 16–48h after transfection. Borosilicate glass pipettes (Harvard Apparatus) pulled to a resistance of 2–6 MΩ (P-1000; Sutter Instrument) and filled with an internal solution containing (in mM): 20 EGTA, 10 HEPES, 50 CsCl, 10 NaCl, and 60 CsF, pH 7.2. Extracellular solution contained (in mM): 110 NaCl, 5 KCl, 40 NMDG, 10 MES, 10 HEPES, 5 glucose, 10 Trizma base, 2 CaCl2, and 1 MgCl2, and pH was adjusted as desired with HCl or NaOH. An Axopatch 200B amplifier and pCLAMP 10.6 (Axon Instruments) were used to record whole-cell currents. Recordings were performed at a holding potential of −80 mV with a 5 kHz low-pass filter and sampling at 10 kHz. Solution changes were performed through rapid perfusion using a SF-77B Fast-Step perfusion system (Warner Instruments). Fluorescence was visualized on an Olympus IX73 microscope (Olympus) with a CoolLED pE-4000 illumination system (CoolLED).

### Preparation and Application of Lipids

Experiments in the presence of PUFA were performed under identical conditions as control experiments except solutions contained the indicated concentration of PUFA. PUFAs stock solutions were made up in ethanol added to aqueous solution following the manufacturer’s recommendations. Ethanol solvent in final solution was typically 0.01% and never exceeded 0.1%. Solution pH was measured prior to and after addition of PUFA to solution to ensure no pH change occurred. Experiments in the presence of PUFA were all performed with at least a 10min preincubation with PUFA to ensure equilibrium was reached. The one exception was for experiments where ASIC currents were monitored on the same cell ± PUFA, a shorter two-minute incubation with PUFA was used. Our time course data suggests that this time period should be sufficient to reach equilibrium.

To determine the pH dependence of activation we measured a series of sweeps from a holding pH of 8 for 7s followed by a 3s activation pulses at indicated pH. Each activation pH was normalized to the maximally activating pH measured at pH 5.5. For pH dependences using rASIC1a, the same protocol was used except the length of the holding pulse was 50 s and the activation pulse was 700ms to minimize the contribution of tachyphylaxis. Each experiment was measured for 3 sweeps and then averaged.

For pH dependences of desensitization, each sweep was performed using a holding pH of 8 for 5s followed by a 22s application of the desensitization solution at the indicated pH. The amount of desensitization was determined at the end of each sweep by measuring the current elicited by a 1.5s pulse of pH 6.0 solution. Each current measured at pH 6.0 was then normalized to the maximal current produced by pH 6.0 determined from a desensitization pH of 8. Each experiment was measured for 3 sweeps and then averaged.

For paired concentration dependences measured at a single pH, controls were measured using a holding pH of 8.0 applied for 90s, followed by 1.5s application of pH 5.5, followed again by 90s at pH 8.0 and finally 3s application of pH 6.6. Following the control recording, an identical protocol was followed for increasing concentrations of PUFA on the same cell. Currents in the presence of PUFA were divided by the pH 6.6 current for their respective controls before averages were calculated.

To measure the I_MAX_ increases, cells in the absence of PUFA were exposed to a holding pH of 8 followed by a 1.5s application of pH 5.5. This was followed by incubation with 10μM PUFA achieved by perfusing 10μM PUFA in pH 8 solution for 2min. The activation protocol was then repeated with 10μM PUFA in both pH 8 and pH 5.5 solutions. Following this, incubation, and activation with 50μM PUFA was performed in an identical manner as 10μM. The pH 5.5 current peaks in the presence of PUFA were divided by the pH 5.5 currents in the absence of PUFA to determine the % I_MAX_. Each experiment was measured for 3 sweeps and then averaged.

To measure desensitization rates, cells were exposed to a holding pH of 8 followed by 3s application of pH 5.5 either with or without 10μM or 50μM PUFA present. For measurement of the AS desensitization rates pH 5.5 was applied for 6s. Rates were determined by taking the time point at which 1/e (63%) of the peak of the pH 5.5 current was reached. Each experiment was measured for 3 sweeps and then averaged.

### Data and Statistical Analysis

PUFA structures and pKa determinations for the various PUFA head group derivatives were made using MarvinSketch chemical editing software (ChemAxon). Calculations of pKa PUFA molecule (head and tail) in the absence of a bilayer at room temperature. Whole-cell patch clamp current recordings were analyzed using Clampfit 10.6 (Axon Instruments). All data are given as mean ± the standard error of the mean (SEM).

For pH dependences, reported pH_0.5_ values represent the mean of the pH_0.5_ values for each individual experiment as determined by fitting to a hill-type equation in SigmaPlot10.0:

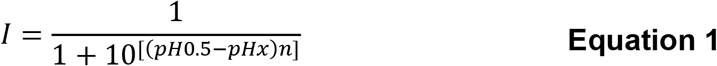

where n = hill number and pH_0.5_=half-maximal activating pH.

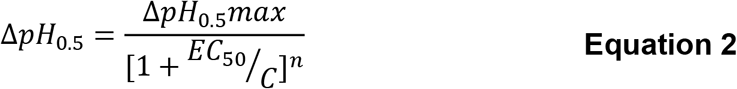

where n = hill number, EC_50_ = half maximal potentiation, and ΔpH_0.5_ = shift in the pH_0.5_.

For wash on kinetics, data was normalized setting the control to zero and the maximum current response obtained during wash on to 1. Reported means for the time to reach 1/e (63%) of the maximal response represent the average value for each individual experiment as determined by fitting to an exponential rise to max in SigmaPlot10.0:

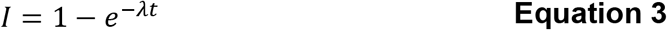

where I = fraction of maximal current, λ = rate constant, and t = time.

For wash off kinetics, data was normalized on the same scale as wash on. Therefore, non-zero minimum values reported for wash off represent the fraction of control prior to wash on. Reported means for the time to wash off 1/e (63%) of the maximal response at time 0s represent the average value for each individual experiment as determined by fitting to an exponential decay in SigmaPlot10.0:

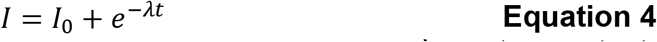

where I = fraction of maximal current, I0 = minimal current response, λ = rate constant, and t = time.

Statistical testing was performed using the program package R. For statistical testing, p-values are reported in tables as calculated except in cases where p-values were <0.001, where they are reported as such. Figure legends indicate which statistical test was used for each data set. For figures, statistical significance is indicated by the following scale: * p < 0.05, ** p <0.01, *** p < 0.001. All p-values reported account for multiple comparisons using appropriate test (see below).

For all statistical testing requiring multiple comparisons, a one-way ANOVA was run followed by post-hoc testing. p-values reported in Table 1 represent a Dunnett’s post-hoc test performed on all pH_0.5_ data sets in the table compared to the control ASIC3 pH_0.5_ dataset. Statistical testing of PUFA effects on desensitization rates and I_MAX_ were performed in a similar manner comparing control ASIC3 datasets to all other datasets using a post-hoc Dunnett’s test. A post-hoc Tukey’s test was also used where indicated to compare between groups means of all data sets.

Paired or unpaired student t-tests were performed where indicated for statistical testing between 2 sets of data (e.g., Supplementary Table 1). All t-tests are two-tailed and unpaired t-tests assume unequal variance.

## Acknowledgements

This work was supported by the National Eye Institute R00 EY024267 (to J.R.B.) and the National Institute of General Medical Sciences R35 GM137912 (to J.R.B.) as well as funding from T32 HL007822 (to R.C.K.).

## Author Contributions

J.R.B. and R.C.K. designed the experiments. R.C.K. performed the experiments. J.R.B. and R.C.K. analyzed the data. J.R.B. and R.C.K. wrote the manuscript.

---

**Supplementary Table 1:** Activation pH_0.5_ values for ASIC3 and ASIC1a mutants. ASIC3 WT data shown again from Table 1 for comparison. p-values for all other constructs were calculated from unpaired t-tests comparing Control to 10μM DHA or AG datasets.

| <u>Construct</u> | <u>Condition</u> | <u>pH<sub>0.5</sub></u> | <u>SEM</u> | <u>n</u> | <u>p-value</u> |
| --- | --- | --- | --- | --- | --- |
| ASIC3 WT | Control | 6.56 | 0.01 | 70 | < 0.001 |
|  | 10μM DHA | 6.83 | 0.03 | 5 |  |
| ASIC3(R64Q) | Control | 6.86 | 0.03 | 10 | 0.593 |
|  | 10μM DHA | 6.85 | 0.02 | 7 |  |
| ASIC3(R64Q) | Control | 6.81 | 0.04 | 4 | 0.020 |
|  | 10μM AG | 7.18 | 0.11 | 4 |  |
| ASIC3(Y67C) | Control | 6.50 | 0.04 | 4 | 0.025 |
|  | 10μM DHA | 6.64 | 0.00 | 4 |  |
| ASIC3(Y68W) | Control | 6.46 | 0.00 | 5 | < 0.001 |
|  | 10μM DHA | 6.57 | 0.02 | 4 |  |
| ASIC3(K428A) | Control | 6.27 | 0.02 | 5 | 0.017 |
|  | 10μM DHA | 6.57 | 0.03 | 4 |  |
| ASIC1a WT | Control | 6.43 | 0.05 | 7 | <0.001 |
|  | 10μM DHA | 6.70 | 0.02 | 6 |  |
| ASIC1a(R64Q) | Control | 6.48 | 0.03 | 8 | 0.082 |
|  | 10μM DHA | 6.58 | 0.04 | 8 |  |
| <u>Non-functional/poorly expressing mutants</u> |  |  |  |  |  |
| <u>Construct</u> | <u>Observable Currents</u> |  |  |  |  |
| ASIC3(R64A) | 0/7 cells |  |  |  |  |
| ASIC3(Y67,68A) | 0/8 cells |  |  |  |  |
| ASIC3(Y431A) | 2/28 cells |  |  |  |  |
| ASIC3(Y431C) | 1/17 cells |  |  |  |  |
| ASIC1a(Y424C) | 0/5 cells |  |  |  |  |
| ASIC1a(Y424W) | 0/7 cells |  |  |  |  |

**Supplementary Table 2:** Abbreviations used throughout the manuscript and calculated pKa values for AA head group derivatives.

| <u>Tail</u> | <u>Abbreviation</u> | <u>Head Group</u> | <u>Calculated $pK_a$</u> |
| --- | --- | --- | --- |
| 22(4,10,13,16) | AA | Carboxyl | 4.83 |
| 22(4,7,10,13,16,19) | DHA | Carboxyl |  |
| 22(4,7,10,13,16) | DPA | Carboxyl |  |
| 20(5,8,11,14) | MA | Methyl Amide | >15 |
| 20(5,8,11,14) | EE | Ethyl Ester | NA |
| 20(5,8,11,14) | E | Ethanolamide | >15 |
| 20(5,8,11,14) | ME | Methyl Ester | NA |
| 20(5,8,11,14) | A | Alanine | 4.02 |
| 20(5,8,11,14) | AS | Serine | 3.81 |
| 20(5,8,11,14) | AG | Glycine | 3.98 |
| 20(5,8,11,14) | EP | Ethanolamide Phosphate | 1.49, 6.49 |
| 20(5,8,11,14) | T | Taurine | -0.83 |

## References

1. Boscardin, E., Alijevic, O., Hummler, E., Frateschi, S. & Kellenberger, S. The function and regulation of acid-sensing ion channels (ASICs) and the epithelial Na+ channel (ENaC): IUPHAR Review 19. Br J Pharmacol 173, 2671–2701 (2016).

2. Dibas, J., Al-Saad, H. & Dibas, A. Basics on the use of acid-sensing ion channels’ inhibitors as therapeutics. Neural Regen Res 14, 395–398 (2019).

3. Du, J. et al. Protons are a neurotransmitter that regulates synaptic plasticity in the lateral amygdala. PNAS 111, 8961–8966 (2014).

4. Wang, X. et al. Serotonin facilitates peripheral pain sensitivity in a manner that depends on the nonproton ligand sensing domain of ASIC3 channel. J Neurosci 33, 4265–4279 (2013).

5. Yu, Y. et al. A Nonproton Ligand Sensor in the Acid-Sensing Ion Channel. Neuron 68, 61–72 (2010).

6. Immke, D. C. & McCleskey, E. W. Lactate enhances the acid-sensing Na+ channel on ischemia-sensing neurons. Nat Neurosci 4, 869–870 (2001).

7. Cadiou, H. et al. Modulation of acid-sensing ion channel activity by nitric oxide. J Neurosci 27, 13251–13260 (2007).

8. Huang, C. L., Feng, S. & Hilgemann, D. W. Direct activation of inward rectifier potassium channels by PIP2 and its stabilization by Gbetagamma. Nature 391, 803–806 (1998).

9. Loussouarn, G. et al. Phosphatidylinositol-4,5-bisphosphate, PIP2, controls KCNQ1/KCNE1 voltage-gated potassium channels: a functional homology between voltage-gated and inward rectifier K+ channels. EMBO J 22, 5412–5421 (2003).

10. Levitan, I., Singh, D. K. & Rosenhouse-Dantsker, A. Cholesterol binding to ion channels. Front. Physiol. 5, (2014).

11. Tian, Y. et al. Atomic determinants of BK channel activation by polyunsaturated fatty acids. Proc Natl Acad Sci U S A 113, 13905–13910 (2016).

12. Hoshi, T., Xu, R., Hou, S., Heinemann, S. H. & Tian, Y. A point mutation in the human Slo1 channel that impairs its sensitivity to omega-3 docosahexaenoic acid. J Gen Physiol 142, 507–522 (2013).

13. Hoshi, T., Tian, Y., Xu, R., Heinemann, S. H. & Hou, S. Mechanism of the modulation of BK potassium channel complexes with different auxiliary subunit compositions by the omega-3 fatty acid DHA. Proc Natl Acad Sci U S A 110, 4822–4827 (2013).

14. Bohannon, B. M. et al. Polyunsaturated fatty acid analogues differentially affect cardiac NaV, CaV, and KV channels through unique mechanisms. eLife 9, e51453 (2020).

15. Liin, S. I., Yazdi, S., Ramentol, R., Barro-Soria, R. & Larsson, H. P. Mechanisms Underlying the Dual Effect of Polyunsaturated Fatty Acid Analogs on Kv7.1. Cell Rep 24, 2908–2918 (2018).

16. Liin, S. I. et al. Polyunsaturated fatty acid analogs act antiarrhythmically on the cardiac IKs channel. Proc Natl Acad Sci U S A 112, 5714–5719 (2015).

17. Bohannon, B. M. et al. Polyunsaturated fatty acids produce a range of activators for heterogeneous IKs channel dysfunction. J Gen Physiol 152, e201912396 (2020).

18. Larsson, J. E., Larsson, H. P. & Liin, S. I. KCNE1 tunes the sensitivity of KV7.1 to polyunsaturated fatty acids by moving turret residues close to the binding site. Elife 7, e37257 (2018).

19. Bohannon, B. M., Perez, M. E., Liin, S. I. & Larsson, H. P. ω-6 and ω-9 polyunsaturated fatty acids with double bonds near the carboxyl head have the highest affinity and largest effects on the cardiac IKs potassium channel. Acta Physiol (Oxf) 225, e13186 (2019).

20. Liin, S. I., Larsson, J. E., Barro-Soria, R., Bentzen, B. H. & Larsson, H. P. Fatty acid analogue N-arachidonoyl taurine restores function of IKs channels with diverse long QT mutations. eLife 5, e20272 (2016).

21. Yoder, N. & Gouaux, E. The His-Gly motif of acid-sensing ion channels resides in a reentrant ‘loop’ implicated in gating and ion selectivity. eLife 9, e56527 (2020).

22. Yoder, N., Yoshioka, C. & Gouaux, E. Gating mechanisms of acid-sensing ion channels. Nature 555, 397–401 (2018).

23. Baconguis, I., Bohlen, C. J., Goehring, A., Julius, D. & Gouaux, E. x-ray structure of acid-sensing ion channel 1-snake toxin complex reveals open state of a Na(+)-selective channel. Cell 156, 717–729 (2014).

24. Jasti, J., Furukawa, H., Gonzales, E. B. & Gouaux, E. Structure of acid-sensing ion channel 1 at 1.9 A resolution and low pH. Nature 449, 316–323 (2007).

25. Allen, N. J. & Attwell, D. Modulation of ASIC channels in rat cerebellar Purkinje neurons by ischaemia-related signals. J Physiol 543, 521–529 (2002).

26. Smith, E. S., Cadiou, H. & McNaughton, P. A. Arachidonic acid potentiates acid-sensing ion channels in rat sensory neurons by a direct action. Neuroscience 145, 686–698 (2007).

27. Deval, E. et al. ASIC3, a sensor of acidic and primary inflammatory pain. EMBO J 27, 3047–3055 (2008).

28. Marra, S. et al. Non-acidic activation of pain-related Acid-Sensing Ion Channel 3 by lipids. EMBO J 35, 414–428 (2016).

29. O’Brien, J. S. & Sampson, E. L. Fatty acid and fatty aldehyde composition of the major brain lipids in normal human gray matter, white matter, and myelin *. Journal of Lipid Research 6, 545–551 (1965).

30. Wiktorowska-Owczarek, A., Berezińska, M. & Nowak, J. Z. PUFAs: Structures, Metabolism and Functions. Adv Clin Exp Med 24, 931–941 (2015).

31. Diochot, S. et al. Black mamba venom peptides target acid-sensing ion channels to abolish pain. Nature 490, 552–555 (2012).

32. Xiong, Z.-G. et al. Neuroprotection in Ischemia: Blocking Calcium-Permeable Acid-Sensing Ion Channels. Cell 118, 687–698 (2004).

33. Basak, S., Schmandt, N., Gicheru, Y. & Chakrapani, S. Crystal structure and dynamics of a lipid-induced potential desensitized-state of a pentameric ligand-gated channel. eLife https://elifesciences.org/articles/23886 (2017) doi:10.7554/eLife.23886.

34. Börjesson, S. I., Hammarström, S. & Elinder, F. Lipoelectric modification of ion channel voltage gating by polyunsaturated fatty acids. Biophys J 95, 2242–2253 (2008).

35. Börjesson, S. I. & Elinder, F. An electrostatic potassium channel opener targeting the final voltage sensor transition. Journal of General Physiology 137, 563–577 (2011).

36. Sherratt, S. C. R. & Mason, R. P. Eicosapentaenoic acid and docosahexaenoic acid have distinct membrane locations and lipid interactions as determined by X-ray diffraction. Chem Phys Lipids 212, 73–79 (2018).

37. Feller, S. E., Gawrisch, K. & MacKerell, A. D. Polyunsaturated Fatty Acids in Lipid Bilayers: Intrinsic and Environmental Contributions to Their Unique Physical Properties. J. Am. Chem. Soc. 124, 318–326 (2002).

38. Varnum, M. D., Black, K. D. & Zagotta, W. N. Molecular mechanism for ligand discrimination of cyclic nucleotide-gated channels. Neuron 15, 619–625 (1995).

39. Wu, Y., Chen, Z., Sigworth, F. J. & Canessa, C. M. Structure and analysis of nanobody binding to the human ASIC1a ion channel. eLife 10, e67115 (2021).

40. Bai, J.-Y., Ding, W.-G., Kojima, A., Seto, T. & Matsuura, H. Putative binding sites for arachidonic acid on the human cardiac Kv1.5 channel. British Journal of Pharmacology 172, 5281–5292 (2015).

41. Chen, Z., Kuenze, G., Meiler, J. & Canessa, C. M. An arginine residue in the outer segment of hASIC1a TM1 affects both proton affinity and channel desensitization. J Gen Physiol 153, (2021).

42. Whitlock, J. M. & Hartzell, H. C. A Pore Idea: the ion conduction pathway of TMEM16/ANO proteins is composed partly of lipid. Pflugers Arch 468, 455–473 (2016).

43. Yazdi, S. et al. Identification of PUFA interaction sites on the cardiac potassium channel KCNQ1. Journal of General Physiology 153, (2021).

44. Farag, N. E., Jeong, D., Claydon, T., Warwicker, J. & Boyett, M. R. Polyunsaturated fatty acids inhibit Kv1.4 by interacting with positively charged extracellular pore residues. Am J Physiol Cell Physiol 311, C255–268 (2016).

45. Paukert, M., Chen, X., Polleichtner, G., Schindelin, H. & Gründer, S. Candidate amino acids involved in H+ gating of acid-sensing ion channel 1a. J Biol Chem 283, 572–581 (2008).

46. Babini, E., Paukert, M., Geisler, H.-S. & Gründer, S. Alternative Splicing and Interaction with Di-and Polyvalent Cations Control the Dynamic Range of Acid-sensing Ion Channel 1 (ASIC1)*. Journal of Biological Chemistry 277, 41597–41603 (2002).

47. Immke, D. C. & McCleskey, E. W. Protons open acid-sensing ion channels by catalyzing relief of Ca2+ blockade. Neuron 37, 75–84 (2003).

48. Paukert, M., Babini, E., Pusch, M. & Gründer, S. Identification of the Ca2+ Blocking Site of Acid-sensing Ion Channel (ASIC) 1. J Gen Physiol 124, 383–394 (2004).

49. Palmer, M. J., Hull, C., Vigh, J. & von Gersdorff, H. Synaptic Cleft Acidification and Modulation of Short-Term Depression by Exocytosed Protons in Retinal Bipolar Cells. J. Neurosci. 23, 11332–11341 (2003).

50. Fraser, D. D. et al. Elevated polyunsaturated fatty acids in blood serum obtained from children on the ketogenic diet. Neurology 60, 1026–1029 (2003).

51. Elinder, F. & Liin, S. I. Actions and mechanisms of polyunsaturated fatty acids on voltage-gated ion channels. Front. Physiol. 8, (2017).

52. Siddiqui, R. A., Harvey, K. A. & Zaloga, G. P. Modulation of enzymatic activities by n-3 polyunsaturated fatty acids to support cardiovascular health. J Nutr Biochem 19, 417–437 (2008).

53. Kuriki, K. et al. Discrepancies in dietary intakes and plasma concentrations of fatty acids according to age among Japanese female dietitians. Eur J Clin Nutr 56, 524–531 (2002).

54. Hammarström, S. et al. Increased concentrations of nonesterified arachidonic acid, 12L-hydroxy-5,8,10,14-eicosatetraenoic acid, prostaglandin E2, and prostaglandin F2alpha in epidermis of psoriasis. PNAS 72, 5130–5134 (1975).

55. Skinner, E. R., Watt, C., Besson, J. A. & Best, P. V. Differences in the fatty acid composition of the grey and white matter of different regions of the brains of patients with Alzheimer’s disease and control subjects. Brain 116 (Pt 3), 717–725 (1993).

56. Anderson, R. E. Lipids of ocular tissues: IV. A comparison of the phospholipids from the retina of six mammalian species. Experimental Eye Research 10, 339–344 (1970).

57. Barbara, G. et al. T-Type Calcium Channel Inhibition Underlies the Analgesic Effects of the Endogenous Lipoamino Acids. J. Neurosci. 29, 13106–13114 (2009).

58. Klipp, R. C., Cullinan, M. M. & Bankston, J. R. Insights into the molecular mechanisms underlying the inhibition of acid-sensing ion channel 3 gating by stomatin. J Gen Physiol 152, e201912471 (2020).

